# DNA methylation profiles in diabetic embryos

**DOI:** 10.1101/2025.10.17.683187

**Authors:** Pu-Yu Li, Fu-Li Zhao, Yi-Juan Song, Bing-Yao Lei, Hao-Jin Niu, Ying-Fang Wang, Hong-Wei Jiang

**Affiliations:** Department of General Medicine, The First Affiliated Hospital of Henan University of Science and Technology, Luoyang, China, 471000; Department of Oncology, The First Affiliated Hospital of Henan University of Science and Technology, Luoyang, China, 471000; Department of Public Health, School of Basic Medical Sciences, Henan University of Science and Technology, Luoyang, China, 471000; Department of Endocrinology, The First Affiliated Hospital of Henan University of Science and Technology, Luoyang, China, 471000

**Keywords:** maternal diabetes, neural tube defects, DNA methylation, Sphk1, Twist1, Shroom3

## Abstract

Maternal diabetes leads to neural tube defects (NTDs) in offspring. DNA methylation could be associated with NTDs induced by maternal diabetes. This study investigated whether maternal diabetes disturbs global DNA methylation of the embryos and possible influence on neural tube closure related genes. Our study showed that there was no significant difference in the overall distribution of DNA methylation levels between nondiabetic and diabetic embryos. The methylation levels of differentially methylated CpGs (dmCpGs) around transcription start sites, gene body and transcription end sites were higher in nondiabetic embryos than that in diabetic embryos. However, there was an increased methylation level of dmCpGs in CpG islands in diabetic embryos compared to that of nondiabetic embryos. Meanwhile, maternal diabetes significantly resulted in differential methylation of CpGs in promoter and gene bodies region of genes such as Sphingosine kinase-1 (Sphk1), twist basic helix-loop-helix transcription factor 1 (Twist1) and shroom family member 3 (Shroom3) and gene expression levels of three genes were altered by maternal diabetes. This study demonstrates that changes of DNA methylation in diabetic embryos led to aberrant expression of genes that may be involved in the pathophysiology of NTDs induced by maternal diabetes.

## Introduction

Maternal diabetes during pregnancy is a well-known teratogen that increases the risk for birth defects, such as neural tube defects (NTDs)^1–3^. The pathophysiology of maternal diabetes-induced birth defects is complicated, however, clearly associated with maternal glucose levels^4^. Although the mechanism is not entirely understood, animal studies have shown that inhibition of cell proliferation and elevation of apoptosis play a vital role on this process^5^. The second major change found on animal studies is altered gene expression that causes deviation from normal developmental process^6–8^. DNA microarray analysis shows significant alterations in expression of developmental and stress response genes in diabetic embryopathy^6–8^. The enrichment of those genes encoding transcriptional regulatory molecules is the most likely candidates to contribute to the molecular etiology for maternal diabetes-induced birth defects^6–8^. However, what causes the alterations of developmental and stress response genes in maternal diabetes conditions is still elusive.

Epigenetic mechanisms, such as DNA methylation, play important roles in etiologies of complex disease such as NTDs^9^. DNA methylation is an important form of epigenetic modification and has a crucial role in many biological processes, including alterations of gene transcription during fetal development^10^. Disruption of embryonic DNA methylation has been experimentally proved to be linked to NTDs^11^. Inactivation of DNA methyltransferase DNMT3B disrupts de novo DNA methylation that causes multiple birth defects including NTDs in mice^12^. It has been reported that maternal nutrition and metabolic disturbances during fetal development can alter epigenetic status such as histone modifications and DNA methylation in fetus^13^. It is well-known that folic acid supplementation prevents NTDs by stimulating cellular methylation reactions^14^. Maternal administration of folic acid diminishes the risk of neural tube defects (NTDs) in offspring^15^. During early embryogenesis, DNA methylation is chief regulator of gene expression^16^. Therefore, we hypothesized that maternal diabetes may alter developmental and stress response genes through the change of DNA methylation that causes the incidence of NTDs.

In the present study, we analyzed the genome-wide DNA methylation pattern in E8.5 embryos (neurulation stage) from maternal diabetic or non-diabetic female mice using reduced representation bisulfate sequencing. We sought to determine whether there are alterations of embryonic DNA methylation pattern in maternal diabetic conditions. Further we assessed whether neural tube closure essential (NTC) genes were differentially methylated between diabetic and nondiabetic embryos. Together, our study provides the detailed map of mouse methylome in neurulation stage embryos (E8.5) and links the change between DNA methylation and NTC genes under maternal diabetic conditions.

## Materials and Methods

### Animals and reagents

Wild-type (WT) C57BL/6J mice were purchased from Jackson Laboratory. Streptozotocin (STZ) from Sigma was dissolved in sterile 0.1 M citrate buffer (pH4.5). The study protocol was approved by the Ethics Committee of The Frist Affiliated Hospital of Henan University of Science and Technology (Approval No. D-2025-B002).

### Mouse models of diabetic embryopathy

The mouse model of diabetic embryopathy has been described previously^5^. Briefly, ten-week-old WT female mice were intravenously injected daily with 75 mg/kg STZ over two days to induce diabetes. Diabetes was defined as 12 hours fasting blood glucose level of ≥ 14 mM. Male and female mice were paired at 3:00 P.M., and pregnancy was established by the presence of the vaginal plug next morning, and noon of that day was designated as day 0.5 (E0.5). On E8.75 (at 6:00 P.M.), mice were euthanized, and conceptuses were dissected out of the uteri, embryos with the yolk sacs were removed from the deciduas and then yolk sacs were removed from the embryos. The embryos from one litter were pooled and three independent biologic replicate samples were obtained for DNA methylation profiles.

### Methyl-MiniSeq™ library construction

Libraries were prepared from 200-500 ng of genomic DNA digested with 60 units of TaqαI and 30 units of MspI (NEB) sequentially and then extracted with Zymo Research (ZR) DNA Clean & Concentrator™-5 kit (Cat#: D4003). Fragments were ligated to pre-annealed adapters containing 5’-methyl-cytosine instead of cytosine according to Illumina’s specified guidelines (www.illumina.com). Adaptor-ligated fragments of 150–250 bp and 250–350 bp in size were recovered from a 2.5% NuSieve 1:1 agarose gel (Zymoclean™ Gel DNA Recovery Kit, ZR Cat#: D4001). The fragments were then bisulfite-treated using the EZ DNA Methylation-Lightning™ Kit (ZR, Cat#: D5020). Preparative-scale PCR was performed, and the resulting products were purified (DNA Clean & Concentrator™ - ZR, Cat#D4005) for sequencing on an Illumina HiSeq.

### Methyl-MiniSeq™ Sequence alignments and data analysis

Sequence reads from bisulfite-treated EpiQuest libraries were identified using standard Illumina base-calling software and then analyzed using a Zymo Research proprietary analysis pipeline, which is written in Python and used Bismark (http://www.bioinformatics.babraham.ac.uk/projects/bismark/) to perform the alignment. Index files were constructed using the bismark genome_preparation command and the entire reference genome. The nondirectional parameter was applied while running Bismark. All other parameters were set to default. Filled-in nucleotides were trimmed off when doing methylation calling. The methylation level (methylation degree) of each sampled cytosine was estimated as the number of reads reporting a C, divided by the total number of reads reporting a C or T. Fisher’s exact test or t-test was performed for each CpG site which has at least five reads coverage, and promoter, gene body and CpG island annotations were added for each CpG included in the comparison.

### RNA extraction and Real-time quantitative PCR (RT-qPCR)

Total RNA was isolated from cells using the Trizol reagent (Thermo Scientific, Waltham, CA) and reverse transcribed using the QuantiTect Reverse Trancription Kit (Qiagen, Valencia, CA). RT-qPCR for Sphk1, Twist1, Shroom3 and β-actin was performed using the Maxima SYBR Green/ROX qPCR Master Mix assay (Thermo Scientific, Waltham, CA). The primers for RT-qPCR are listed in Table 1. RT-qPCR and subsequent calculations were performed by a StepOnePlus™ Real-Time PCR System (Applied Biosystem, Foster City, CA).

**Table 1.**
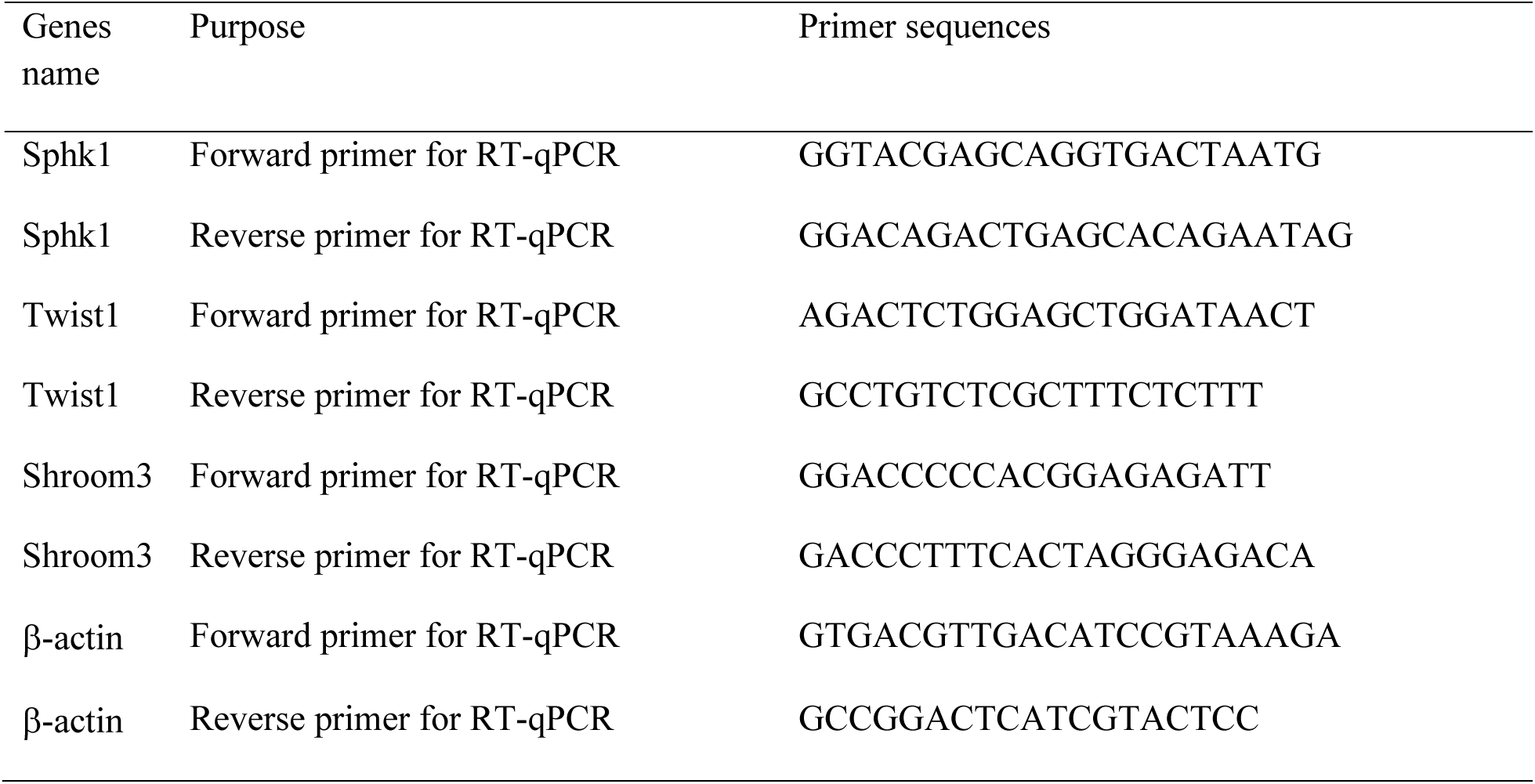
Primers for RT-qPCR.

## Statistical analysis

Data are presented as means ± SEM. Embryonic samples from each replicate were from different dams. Statistical differences were determined by one-way analysis of variance (ANOVA) using SigmaStat 3.5 software (Systat Software Inc., San Jose, CA). In one-way analysis of variance, *Tukey test* was used to estimate the significance of the results (*P* < 0.05).

## Results

### Global DNA methylation characterization in nondiabetic and diabetic embryos

To investigate global DNA methylation changes in E8.5 embryos between diabetic and nondiabetic groups, Methyl-MiniSeq™ (Zymo Research) platform, which is an improved version of Reduced Representation Bisulfite Sequencing (RRBS), was used to map the DNA CpG methylation state on a global scale. The bisulfite conversion rate was almost 100% (99%). For each sample, we obtained approximately 27-28 million reads, and 4 million CpGs were observed and aligned to the mouse genome, mapping about 2.7 million unique CpGs with average 14-fold coverage. The whole sequencing results were visualized with Integrative Genomics Viewer or the UCSC genome browser and can be observed using the supplied kinks of genome browser tracks. Further analysis of those CpGs indicated all the chromosomes were covered by those CpGs with almost equal number, except chromosome Y. 34% of CpGs are located in CpG islands. Meanwhile, 34 % of CpGs are located in promoter region, 59% in gene body (24% in exon, 35% in intron). 33% of CpGs are located in intergenic region.

To describe global differences in embryos between nondiabetic and diabetic groups, we calculated the average level (defined as the ratio of methylated reads/total reads at a given site) of DNA methylation in all CpG sites. Firstly, there was no significant difference in the overall distribution of DNA methylation level between nondiabetic and diabetic embryos (Figure 1A and 1C). Secondly, each chromosome had different DNA methylation level despite no diversity between nondiabetic and diabetic embryos (Figure 1B). Thirdly, we observed the obvious low DNA methylation level in CpG islands, promoter and exon regions in nondiabetic or diabetic embryos (Figure 1C). However, there was a high DNA methylation level in non CpG islands, intron, and intergenic regions with no significant difference between nondiabetic and diabetic embryos (Figure 1C). Fourthly, the DNA methylation level of the gene body (the genomic region from transcription start sites (TSS) of a gene to its transcription end sites (TES) is higher than that of neighboring intergenic regions and there is a markedly hypomethylated region around the TSS (Figure 1D). The methylation level of the gene body is a slight increase from the TSS to the TES and a clear reduction after the TES (Figure 1D). But the pattern is similar between nondiabetic and diabetic embryos.

**Figure 1.**
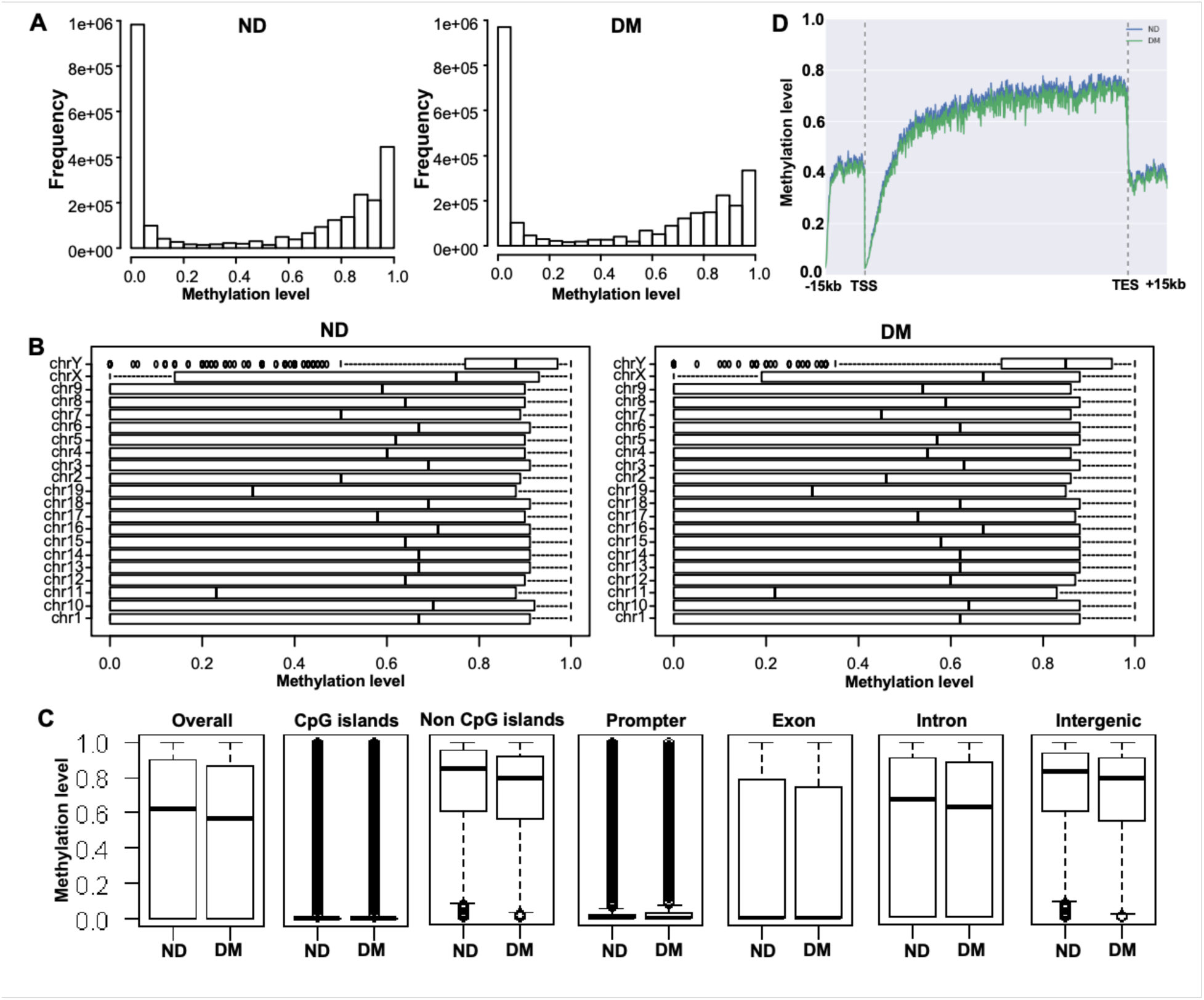
Global DNA methylation level in nondiabetic and diabetic embryos. A. Frequency of DNA methylation level in different degree. B. DNA methylation level in each chromosome. C. Overall DNA methylation level and methylation patterns within different genomic regions including CpGs islands, non CpGs islands, promoter, exon, intron and intergenic regions. D. DNA methylation level along the gene bodies and 15 kilobases (kb) upstream of the transcription start sites (TSS) and 15 kb downstream of the transcription end sites (TES) of all RefSeq genes.

### Genomic distribution of differentially methylated CpGs (dmCpGs) in nondiabetic and diabetic embryos

In the present study, we identified significantly differentially methylated CpGs (dmCpGs) that either gained or lost methylation in diabetic embryos compared to nondiabetic embryos. In total, approximately 53101 (2%) statistically significant dmCpGs (p < 0.05) were identified in diabetic embryos (Figure 2A). Among those dmCpGs, a total of 14963 (0.56%) CpGs had gained methylation (hypermethylation, 0.26% with strong hypermethylation), while 38138 (1.42%) CpGs had lost methylation (hypomethylation, 0.79% with strong hypomethylation) (Figure 2A). Meanwhile, these dmCpGs including hypermethylation and hypomethylation did not accumulate in certain chromosomes. They were distributed in all chromosomes equally, except Y chromosome (Figure 2B). But the number of hypomethylated dmCpGs was higher than that of hypermthylated dmCpGs (Figure 2B). We next compared the genomic distribution of dmCpGs at CpG islands, non CpG islands, promoter, exon, and intron regions of genome between nondiabetic and diabetic embryos. Our results showed that the majority of dmCpGs located in non-CpG islands (Figure 2C and 2D). Meanwhile, hypermethylated CpGs distributed in promoter, exon, intron, and intergenic regions almost equally, but approximately half of hypomethylated CpGs located in intergenic regions (Figure 2E and 2F).

**Figure 2.**
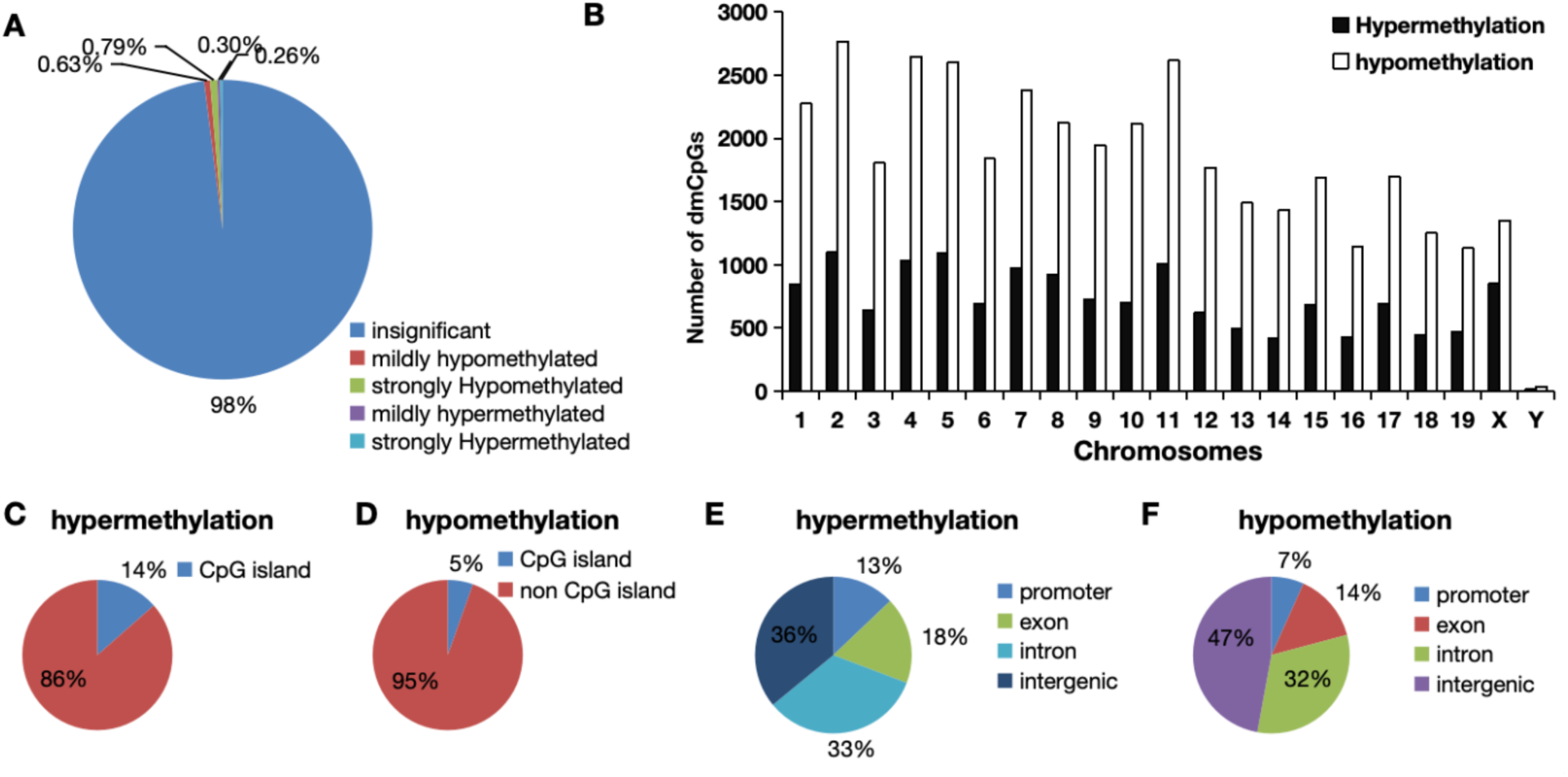
Genomic distribution of differentially methylated CpGs (dmCpGs) in nondiabetic and diabetic embryos. A. Percentage of dmCpGs wih hypermethylation, hypomethylation or insignificant methylation. B. Chromosomal distribution of dmCpGs with hypermethylation or hypomethylation. C. Percentage of dmCpGs with hypermethylation in CpG islands and non CpG islands. D. Percentage of dmCpGs with hypomethylation in CpG islands and non CpG islands. E.Genomic location of dmCpGs with hypermethylation. F. Genomic location of dmCpGs with hypomethylation.

### Methylation level of dmCpGs in different genomic elements in nondiabetic and diabetic embryos

To test whether there is a total difference of dmCpGs in different genomic elements, we calculated methylation level of dmCpGs in diabetic or nondiabetic embryos. First, of those dmCpGs, most of the methylation changes (delta methylation level) ranging from 10% to 50% (Figure 3A). Meanwhile, the methylation level of dmCpGs was similar in each chromosome, except chromosome X (Figure 3B). It seems that methylation level of dmCpGs in nondiabetic embryos was higher than that in diabetic embryos (Figure 3B and 3C). The increase of methylation level of dmCpGs in nondiabetic embryos may result from the increase in exon, intron, and intergenic regions and at the same time those dmCpGs were also located in non CpG islands (Figure 3C and 3D). Although the methylation level of dmCpGs in promoter regions was similar, there was an elevation in CpG islands in diabetic embryos compared to nondiabetic groups (Figure 3C and 3D). However, the methylation level of dmCpGs around TSS, gene body and TES was higher in nondiabetic embryos than that in diabetic embryos even with robust hypomethylation around TSS (Figure 3D).

**Figure 3.**
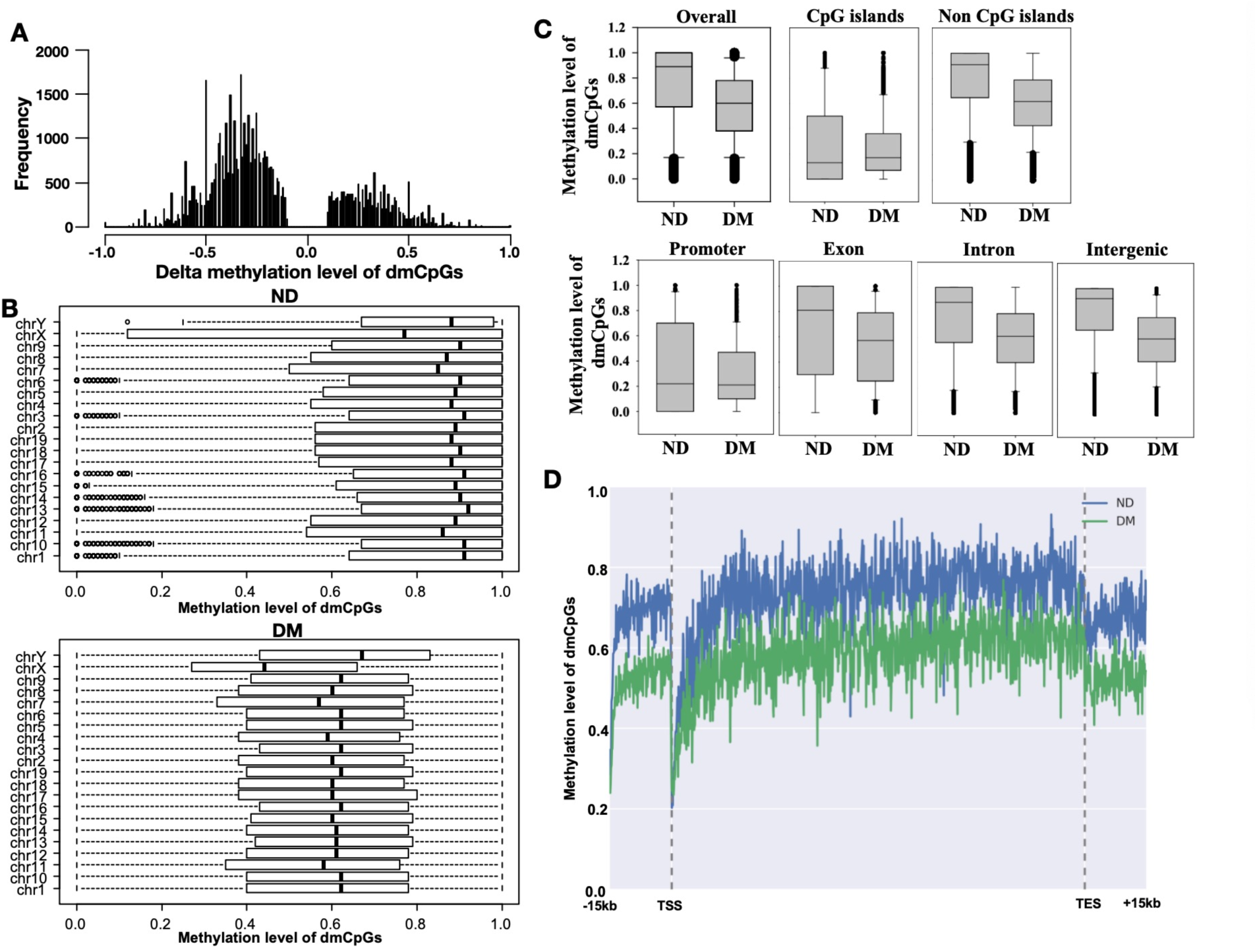
Methylation level of dmCpGs in nondiabetic and diabetic embryos. A. Frequency of delta methylation level of dmCpGs in different degree. Plus (+) means hypermethylation, minus (-) means hypomethylation in diabetic emrbyos. B. Methylation level of dmCpGs in each chromosome. C. Overall methylation level of dmCpGs and methylation patterns within different genomic regions including CpGs islands, non CpG islands, promoter, exon, intron and intergenic regions. D. Methylation level of dmCpGs along the gene bodies and 15 kilobases (kb) upstream of the transcription start sites (TSS) and 15 kb downstream of the transcription end sites (TES) of all RefSeq genes. dmCpGs: differentially methylated CpGs.

### Maternal diabetes induced the change of methylation status in neural tube closure essential genes

Maternal diabetes-induced NTDs are profoundly associated with altered gene expression in developing embryos^6^. These genes were named neural tube closure essential genes (NTC genes), also named NTD-associated genes, the number of which has been identified more than 300 genes^6^.

Salbaum et al obtained 372 of NTC genes by queries of Mouse Genome Informatics (MGI) database and the paper reviewed by Harris and Juriloff ^6,18^. Therefore, we evaluated methylation level of NTC genes in diabetic embryos in the present study. Totally, 206 of 372 NTC genes were significantly hypermethylated or hypomethylated in promoter, exon, or intron under maternal diabetic conditions. Figure 4A and 4B showed the number of NTC genes with strong hypermethylation or hypomethylation (fold change > 30%) in different genomic regions.

**Figure 4.**
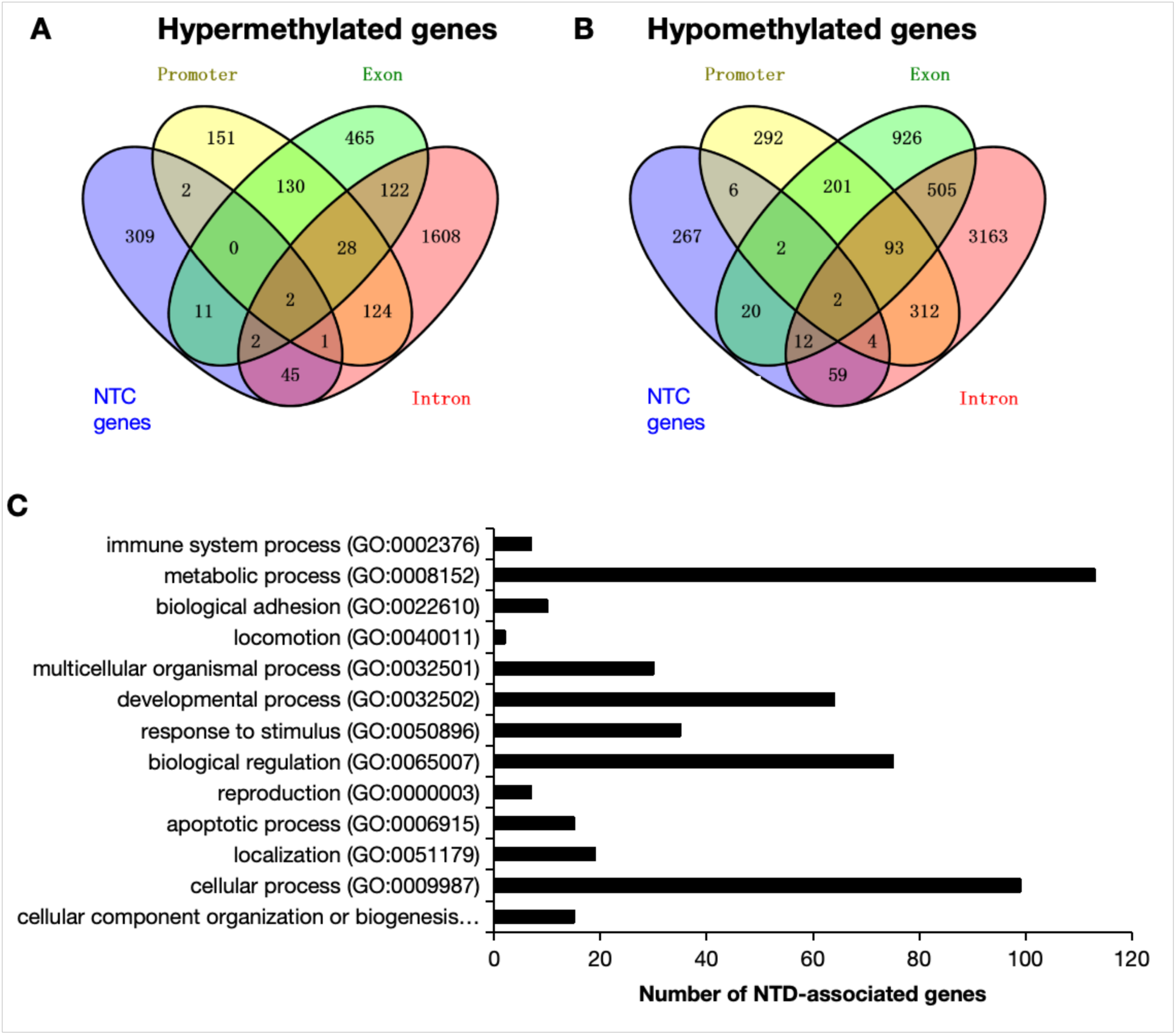
The genomic distribution of dmCpGs and functional annotation of NTC genes. A and B. Venn diagram illustrated genomic location of dmCpGs in promoter, exon or intron of NTC genes. Red: the number of dmCpGs in intron region of genes; Green: the number of dmCpGs in exon region of genes; Yellow: the number of dmCpGs in promoter region of genes; Blue: the number of dmCpGs in NTC genes. Overlapped color indicates the number of dmCpGs which co-located in two or more different genomic elements or NTC genes. C. Functional annotation of NTC genes whose methylation status was affected by maternal diabetes. Category names are presented on the y axis. The x axis indicates the enrichment gene number of NTC genes. The enchrichment of gene number corresponding to each term provides by the Protein Analysis Through Evolutionary Relationships (PANTHER, http://pantherdb.org/). NTC gene: neural tube closure essential genes. dmCpGs: differentially methylated CpGs.

Those differential methylated NTC genes (total 206 genes) were then analyzed in the context of GO analysis based on the biological process using the Protein Analysis Through Evolutionary Relationships (PANTHER) classification system. As shown in Figure 4C, those genes were divided into 13 categories of biological process, 113 of genes in metabolic process and 99 of genes in cellular process. In addition, we found there were 15 of genes in apoptosis process, which were consistent with previous studies that apoptosis is an important causal factor for maternal diabetes-induced NTDs^4^. Taken together, it is suggested that maternal diabetes-induced NTDs may be associated with aberrant methylation in known NTC genes.

### DNA methylation correlates with aberrant expression of NTC genes in diabetic embryos

Three NTC genes (Sphingosine kinase-1 (Sphk1), twist basic helix-loop-helix transcription factor 1 (Twist1) and shroom family member 3 (Shroom3)) were chosen for validation of methylation status and their relationship with gene expression because these NTC genes had more than 7 of dmCpGs in their promoter (Sphk1 and Twist1) or gene body (Shroom3) as well as their dmCpGs belong to CpG islands. Detailed analysis of the DNA methylation pattern in Sphk1, Twist1, and Shroom3 genes is shown in Figure 5A, 5B and 5C using UCSC genome browser tracks to display methylation data at the nucleotide resolution. Sphk1 showed strong hypermethylation in promoter region, in contrast, Twist1 showed strong hypomethylation in promoter region (Figure 5A and 5B). However, there was strong hypomethylation in fifth exon of Shroom3 (Figure 5C). RT-qPCR results indicated the aberrant expression of these genes (Figure 5D). Hypermethylation of promoter leads to decreased expression of genes such as Sphk1 (Figure 5D). In contrast, hypomethylation of promoter results in increased expression of genes such as Twist1 (Figure 5D). Interestingly, hypomethylation of exon triggered the decreased expression of Shroom3 (Figure 5D).

**Figure 5.**
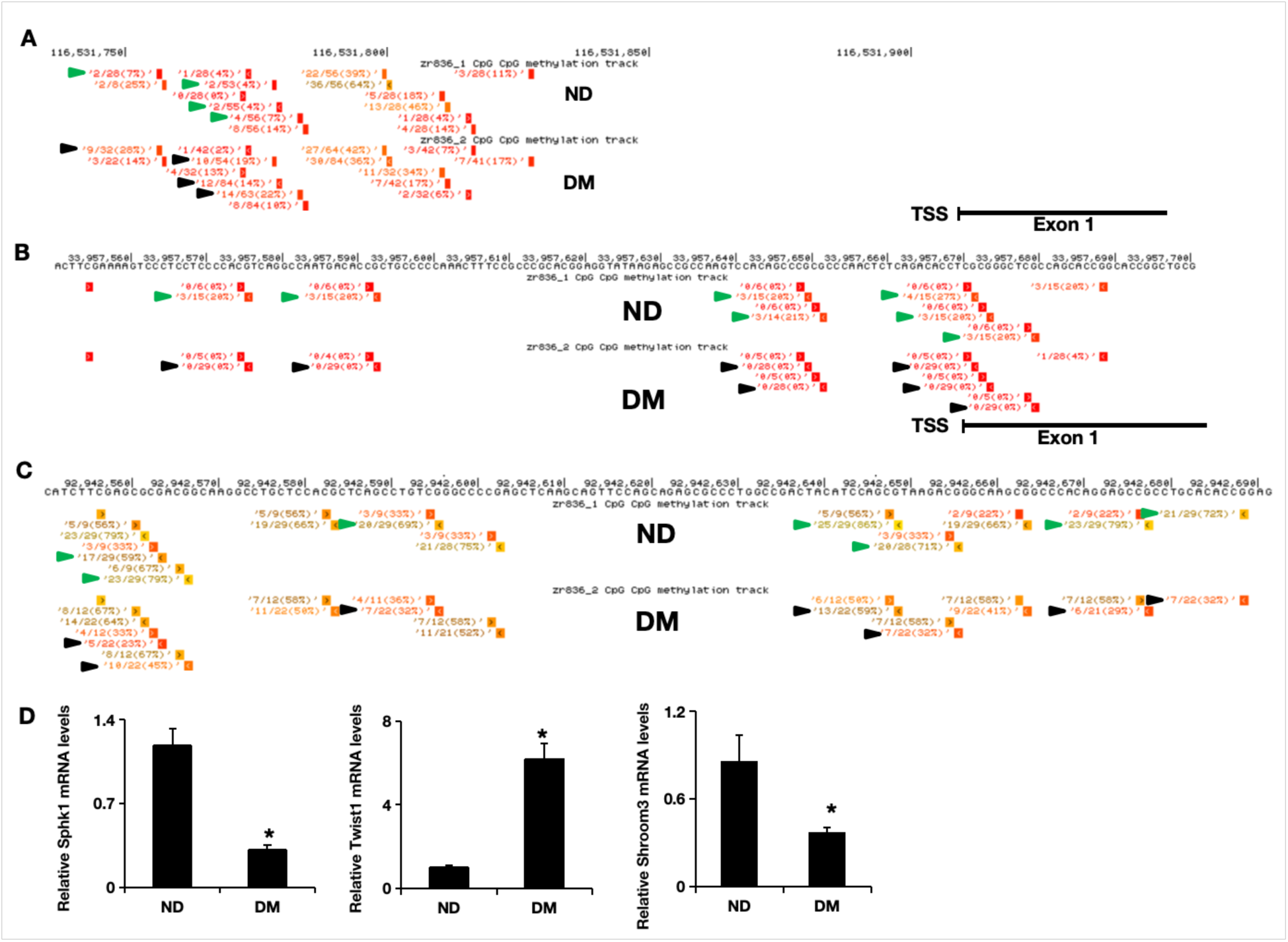
DNA methylation pattern and gene expression of Sphk1, Twist1, Shroom3 in nondiabetic and diabetic embryos. A. The promoter DNA methylation pattern of Sphk1 in nondiabetic and diabetic embryos. B. The promoter DNA methylation pattern of Twist1 in nondiabetic and diabetic embryos. C. The fifth Exon DNA methylation pattern of Shroom3 in nondiabetic and diabetic embryos. DNA methylation data display ratios with the number of methylated reads divided the total number of reads at each indivdual CpG site. The data were visualized using the UCSC genome browser track. High methylation is denoted in yellow and low methylation in red. Green and black arrows indicate the position of dmCpGs. dmCpGs: differentially methylated CpGs. ND: nondiabetic group, DM: diabetic group. D. mRNA levels of Sphk1, Twist1, Shroom3 in nondiabetic (ND) and diabetic (DM) embryos determined by RT-qPCR. Values are means ± SEM from three separate experiments. * indicates significant difference (*P* < 0.05) compared to ND group.

## Discussion

Diabetic mice were defined as blood glucose levels exceeding 16.7 mM. Control mice were maintained at normal blood glucose levels (5-7 mM).The developmental stage E8.5 was chosen for this study because previous studies showed that this stage is important for neurulation and neural tube closure^5^. Therefore, global DNA methylation profiles were performed using E8.5 emrbyos from nondiabetic or diabetic dams. In this study, we determined global DNA methylation profiles and identified 53101 dmCpGs and these dmCpGs exhibited average hypomethylation in diabetic embryos but with an elevation in CpG islands. Meanwhile, we found 206 of NTC genes were differential methylated by maternal diabetes in different genomic elements. Furthermore, these differential methylated NTC genes were categorized according to biological processes and 15 NTC genes were candidate genes for regulation of apoptosis that was demonstrated by previous studies that apoptosis is causative events for maternal diabetes-induced NTDs^4^. The expression of several NTC genes were validated by using RT-qPCR. Here, we provide evidence that maternal diabetes led to the alterations of global DNA methylation pattern that causes aberrant methylation and expression of NTC genes in diabetic embryos.

For genome-wide analysis of DNA methylation changes in nondiabetic and diabetic embryos, an improved version of Reduced Representation Bisulfite Sequencing (BBRS) using next-generation sequencing was applied^19^. BBRS, like some of the other methylation techniques, uses sodium bisulfite modified DNA that is digested with specific restriction endonucleases and enrichment for GC rich regions allowing a focused sequencing of primarily CpG rich genomic regions. It provides a coverage of up to 90% of functionally relevant annotated regions such as CpG islands, promoter and gene bodies, which are of special important with epigenetic gene regulation, while genomic coverage is only around 10-20%^20^. Therefore, our data showed the overall changes of DNA methylation in CpG enrichment regions that is useful for the analysis of maternal diabetes-induced alterations of NTC genes. However, previous data showed overall DNA hypermethylation in diabetic embryos by using DNA methylation ELISA kit^21^. The inconsistency may be attributed to low coverage of non CpG enrichment regions in the BBRS method.

DNA methylation is primarily confined to cytosine bases and is associated with transcriptional silencing^22^. The present of 5-methylcytosine (5-mC) in many promoter causes gene silencing^22^. Our present study showed maternal diabetes induced a lot of dmCpGs in the promoter of NTC genes and hence control their expression pattern. Our results confirmed that hypermethylation in the promoter of genes reduces gene expression, whereas hypomethylation in the promoter of genes elevates gene expression, for example, Sphk1 and Twist1 genes. Meanwhile, a variety of dmCpGs located in gene body (intron and exon) of NTC genes were found in diabetic embryos. The function of DNA methylation in gene body is still elusive and controversial^22^. At least, gene body methylation is not associated with repression because early research demonstrated that gene body methylation is a feature of transcribed genes although some reports showed the opposite results^23,24^. Mechanistic research demonstrated that DNA methylation of gene body stimulates transcription elongation but not initiation^23^. Our results showed that some genes have hypermethylation in gene bodies in nondiabetic embryos with higher gene expression than diabetic embryos, for example, Shroom3. Thus, diverse locations of dmCpGs in genes will lead to different gene expression outcomes and ultimately cause dysregulation of NTC genes in diabetic embryos.

Interestingly, DNA methylation levels of dmCpGs in CpG islands were increased despite there being an overall hypomethylation of dmCpGs and the similar methylation levels of dmCpGs in the promoter in diabetic embryos. CpG islands are defined with higher CpG density than the rest of genome on 1000 base pairs length^24,25^. Approximately 70% of annotated gene promoters are associated with a CpG islands, making this the most common promoter type in the vertebrate genome^25^. Methylation of CpG islands in the promoter can impair transcription factor binding, recruit repressive methyl-binding proteins, and finally stably silence gene expression^24^. Our analysis indicated that more than half of dmCpG located in CpGs islands as well as in the promoter of genes with hypermethylated status under maternal diabetes conditions. The result that there are the similar mentylation levels of promoter between nondiabetic and biabetic embryos is because some dmCpGs in the promoter did not belong to the CpG islands. Thus, it is suggested that maternal diabetes induces the hypermethylation of CpG islands in the promoter of genes that causes the repression of most of genes in developed embryos.

Although it has been widely shown that maternal diabetes-induced NTDs are associated with epigenetic modifications^10^, the knowledge about DNA methylation changes in diabetic embryos is still limited. Several studies investigated alterations of DNA methylation patterns in umbilical cord blood and placenta exposed to maternal diabetes or obesity^26–30^. Spina bifida (one type of NTD) has been thought to be associated with altered pattern of DNA methylation in placenta^31^. At the same time, previous studies showed that maternal diabetes or obesity cause changes of DNA methylation of individual genes in offspring, such as Pax3 with hypermethylation^32^, Peg3 and H19 with hypermethylation^33,34^, GNAS with hypermethylation^35^. In the present study, although we found that methylation levels of dmCpGs were lower in diabetic embryos than that in non-diabetic embryos and global DNA hypomethylation induced by methptrexate inhibition is indeed associated with NTDs^36^, a lot of NTC genes were also hypermethylated at promoter or gene body regions such as Sphk1, Hic1 and Pax5. In future study, we need to unlock possible mechanisms for global hypomethylation and hypermethylation of individual genes in diabetic embryopathy.

GO annotation of differential methylated NTC genes showed enrichment in GO terms, which were associated with biological processes like metabolic, cellular, developmental processes that are important events during neural tube closure. Interestingly, about 15 of neural tube closure essential genes are associated with apoptotic processes such as Sphk1. Sphk1 can enhance resistance to apoptosis through activation of signal pathways in different types of cancer^37–39^. Previous studies have revealed that ER stress^40–43^, oxidative stress^41–44^, suppressed autophagy^45^ and DNA damage^46^ induced apoptosis in diabetic embryos. In the present study, our data demonstrated that aberrance of DNA methylation causes the dysregulation of apoptosis-related genes such as Sphk1 that may be another causative factor for maternal diabetes-induced apoptosis and ultimate NTDs.

Investigations of DNA methylation on pregnancies have important clinical significance. Alteration of DNA methylation on pregnancies have been associated with a variety of adverse pregnancy outcomes including fetal autism spectrum disorders^47^, preterm birth and infection^48^, placental dysfunction^49–51^, Fetal programming and systemic sclerosis^52^, cervical carcinoma^53^, ovarian serous neoplasms^54^ and congenital cardiac malformations^55^. DNA methylation markers were used for non-invasive prenatal diagnosis of aneuploidies^56^ and fetal trisomy 21^57^. Previous study showed that the Green tea polyphenol EGCG supplementation alleviates maternal diabetes– induced NTDs by regulating fetal DNA methylation^21^. The preventive effect of folic acid supplementation in early pregnancy on fetal NTDs is associated with fetal DNA methylation pattern^58–60^. In summary, elucidating DNA methylation alteration in embryos affected by maternal diabetes will significantly advance our understanding of the mechanism of maternal diabetic embropathy. Moreover, defining the molecular targets that lead to DNA methylation alterations and aberrance of NTC genes can potentially be used for designing future clinical applications for treatment of NTDs.

## Acknowledgements

This research was funded by the Luoyang Science and Technology Development Plan Project (2302013Y), Key scientific research projects of colleges and universities in Henan Province (22A310013)

